# Automated hyperspectral vegetation index derivation using a hyperparameter optimization framework for high-throughput plant phenotyping

**DOI:** 10.1101/2021.10.27.466056

**Authors:** Joshua C.O. Koh, Bikram P. Banerjee, German Spangenberg, Surya Kant

## Abstract

- Hyperspectral vegetation indices (VIs) are widely deployed in agriculture remote sensing and plant phenotyping to estimate plant biophysical and biochemical traits. However, existing VIs consist mainly of simple 2-band indices which limits the net performance and often do not generalize well for other traits than they were originally designed for.
- We present an automated hyperspectral vegetation index (AutoVI) system for the rapid generation of novel 2- to 6-band trait-specific indices in a streamlined process covering model selection, optimization and evaluation driven by the tree parzen estimator algorithm. Its performance was tested in generating novel indices to estimate chlorophyll and sugar contents in wheat.
- Results show that AutoVI can rapidly generate complex novel VIs (≥4-band index) which correlated strongly (R^2^ > 0.8) with measured chlorophyll and sugar contents in wheat. AutoVI-derived indices were used as features in simple and stepwise multiple linear regression for chlorophyll and sugar content estimation, and outperformed results achieved with existing 47 VIs and those provided by partial least squares regression.
- The AutoVI system can deliver novel trait-specific VIs readily adoptable in high-throughput plant phenotyping platforms and should appeal to plant scientists and breeders. A graphical user interface of AutoVI is herein provided.

## Introduction

High-throughput plant phenotyping (HTP) is integral in meeting the demand for large-scale evaluation of genotypes in breeding programs and crop management systems (Tardieu *et al*., 2017; Mir *et al*., 2019). In recent years, controlled environment and field-based HTP platforms have been developed to monitor plants at the canopy or plot level for a large number of crop genotypes (Tardieu *et al*., 2017; Mir *et al*., 2019; Lu *et al*., 2020). Central to the success of these HTP platforms is the use of various imaging sensors to acquire morphological, physiological, and biochemical parameters in a non-invasive manner. Hyperspectral imaging has been a promising HTP technology in measuring biochemical and morpho-physiological traits in a fast and non-destructive way by detecting signatures in the reflectance spectrum of vegetation in narrow (e.g. 1 - 2 nm in spectral resolution) and contiguous/broad (e.g. ≥ 20 nm in spectral resolution) spectral bands (Lu *et al*., 2020). The recent availability of lightweight hyperspectral sensors has stimulated a rapid adoption of these sensors for use in unmanned aerial vehicle systems for HTP and precision agriculture (Adão *et al*., 2017; Lu *et al*., 2020). Hyperspectral data have been applied for the estimation of biophysical (e.g. leaf area index, green canopy cover) and biochemical (e.g. chlorophyll, nitrogen, sugar) traits for the leaf and canopy, as well as traits relating to the whole plant, particularly those linked with growth (e.g. plant biomass, plant height) and productivity (e.g. grain yield) (Adão *et al*., 2017; Lu *et al*., 2020). Apart from this, close-range hyperspectral imaging (ground-based or glasshouse) characterized by high spatial resolution and signal-to-noise ratio is also increasingly common in HTP facilities. These systems allow fine-scale investigation of vegetative features at the leaf or canopy level with applications in plant water content and biochemical compounds estimation, and detection of abiotic and biotic stresses in plants (Mishra *et al*., 2017).

However, the large amount of data collected by hyperspectral sensors poses challenges in analytical implementations. Redundancy problems linked to the multicollinearity of bands and the curse of dimensionality impose high computational costs on analytical pipelines (Bajwa & Kulkarni, 2011; Burger & Gowen, 2011). Multicollinearity happens when independent variables (e.g. wavebands) are highly correlated, leading to inaccurate estimation of coefficients (effect of independent variables on response/target trait) in a regression analysis and less reliable statistical inferences (Bajwa & Kulkarni, 2011; Burger & Gowen, 2011). On the other hand, curse of dimensionality refers to the problem of optimizing a given function or model due to the exponential increase in possible solutions or sets of parameters associated with the increase in the number of variables (i.e. the dimensionality), which often necessitates exhaustive enumeration of the solution/search space to achieve satisfactory optimization (Piatetsky-Shapiro *et al*., 2000). Dedicated efforts are often required to develop efficient hyperspectral data processing workflows for a specific plant phenotyping task (Aasen *et al*., 2014; Aasen *et al*., 2018). To this end, vegetation indices (VIs) offer an alternate approach to hyperspectral data analysis which is simple and fast. The VIs are formulated as ratios or algebraic combinations of vegetative reflectance at different wavebands, selected from the visible (VIS, 400 – 700 nm), near infrared (NIR, 700 – 1000 nm) and the shortwave infrared (SWIR, 1000 – 2500 nm) regions (Silleos *et al*., 2006; Xue & Su, 2017). The application of VIs for plant phenotyping is straightforward, requiring the user to compute index values directly from the relevant wavebands’ reflectance and use them either as proxy measures to the target trait or as features (or variables) in regression modelling to predict the target trait values. Since the introduction of the first VI by Pearson and Miller (1972), more than 500 hyperspectral VIs have been developed, demonstrating a strong and continued interest in the development and adoption of novel VIs for specific remote sensing applications (Henrich *et al*., 2017). However, existing VIs consist predominantly of 2-band indices and to some extent, 3-band indices, which limits the amount of information represented and thereby the net performance afforded by these VIs (Henrich *et al*., 2017). In addition, existing VIs are designed for specific plant traits and often do not generalize well for other traits. As such, particularly in agriculture, there is a continued demand for novel VIs that can target specific traits associated with crop growth, biochemical parameters, yield, and quality.

The development of VIs is technically challenging and time-consuming, often requiring a comprehensive understanding of the dynamic changes of the plant optical properties in relation to the intrinsic biochemical or biophysical trait(s) of interest. To this end, a wide variety of experiments have been established to acquire a comprehensive spectral library (Rao *et al*., 2007; Chauhan & Mohan, 2013). Ideally, knowledge on wavebands associated with plant traits may be enriched or expanded with successive development of new VIs when different regions of the reflectance spectrum corresponding to vegetative features are identified. Biochemical and biophysical traits can be described comprehensively with more wavebands where each waveband adds supplementary information. However, this is seldom the case as the VIs rarely constitute a complex or cohesive assemblage of wavebands but rather a limited selection of a few wavebands (usually ≤ 4 bands). In addition, multicollinearity of bands and the curse of high dimensionality inherent in hyperspectral data complicates the identification of wavebands linked to the underlying trait of interest. Several attempts have been made to accelerate the development of novel VIs, these include the use of correlation matrices between VIs and the target traits of interest to retrieve new waveband or index combinations (Thenkabail *et al*., 2004; Aasen *et al*., 2014; Xu *et al*., 2019). Careful selection of hyperspectral features (wavebands) was effective in overcoming the curse of dimensionality and the selected bands could be combined to develop VIs (Aasen et al., 2014; Aasen et al., 2018). Recently, a brute force indices mining approach was applied in our lab to identify a new normalized difference chlorophyll index (NDCI_w_) for the estimation of chlorophyll content in wheat (Banerjee *et al*., 2020). However, these approaches are suited for the evaluation of a limited number of wavebands and/or index model combinations for the development of new VIs; and may be computationally less efficient when dealing with a greater number of wavebands and index models. An efficient VIs evaluation and waveband selection strategy is crucial to developing trait-specific hyperspectral VIs for HTP and agriculture remote sensing.

This study aims to report the development and evaluation of an automated hyperspectral VI (AutoVI) derivation system for the rapid generation of trait-specific novel 2-band to 6-band VIs based on a hyperparameter optimization framework. The term “hyperparameter optimization” (HPO) is often associated with the machine learning discipline in which specific algorithms are deployed to select optimum values in a defined search space for model parameters (values learned from data) and hyperparameters (values associated with the model function or architecture) to maximize the model performance (Bergstra *et al*., 2011; Yu & Zhu, 2020). Building upon an HPO framework, AutoVI is designed to deliver an end-to-end VI development pipeline covering index model evaluation and optimum waveband selection with minimal user input. In this study, novel VIs for chlorophyll and sugar estimation in wheat were generated using AutoVI and compared against existing VIs as features in simple and multiple regression modelling, with the results also compared against those computed by partial least squares regression. The potential application of AutoVI for HTP and agriculture remote sensing is discussed.

## Materials and Methods

### Experiment in a high-throughput phenotyping facility

Data used in this study was collected in a high-throughput controlled-environment phenotyping facility in the Plant Phenomics Victoria, Horsham (PPVH), Agriculture Victoria as previously described (Banerjee *et al*., 2020). In brief, the PPVH facility is equipped with a conveyor belt system, automated weighing and watering stations, and an automated phenotyping Scanalyzer 3D system (LemnatecGmBH, Aachen, Germany), which includes a hyperspectral imaging sensor. For the experiment conducted at the PPVH, wheat plants were grown at 2, 5, 10, and 20 mM nitrogen (N) levels. One plant per pot was grown in a nutrient-free growth medium, consisting of perlite covered with a layer of vermiculite. Individual pots were weighed and equalized to a fixed pot weight and watered uniformly. The pots were loaded onto the system 10 days after the emergence of seedlings. Nutrient solution (4 mM MgSO_4_, 4 mM KCl, 5 mM CaCl_2_, 3 mM KH_2_PO_4_/K_2_HPO_4_-pH 6.0, 0.1 mM Fe-EDTA, 10 μM MnCl_2_, 10 μM ZnSO_4_, 2 μM CuSO_4_, 50 μM H_2_BO_3_, and 0.2 μM Na_2_MoO_4_) with the indicated N concentration was supplied as 100 ml per pot every week. The growing conditions were 16 h (24 °C) day / 8 h (15 °C) night. The experiment was conducted as biological repeats with 20 replicate plants per N treatment. A subset of five plants per N treatment were destructively harvested at 14, 21, 28 and 35 days after sowing (DAS) and a total of 80 samples were collected for biochemical assays.

### Hyperspectral image acquisition and processing

Plants were scanned in an imaging station with a push broom hyperspectral sensor (Micro-Hyperspec, VNIR-E Series, Headwall Photonics, Fitchburg, MA, USA) over a spectral range of 475 – 1710 nm in three viewing angles (0°, 120°, and 240°). Raw data acquired in 12-bit digital numbers (DNs) were transformed to radiance following spectral and radiometric calibration using a white reference target (Spectralon panel, Labsphere Inc, North Sutton, US) and dark reference (spectrum collected with halogen lamps turned off) in a data-acquisition software (Hyperspec III, Headwall Photonics, Inc, Massachusetts, US). Further processing was applied to remove inter-channel variation and correct illumination variations (Banerjee *et al*., 2020). Non-plant pixels (cage, pot, soil, and background) in the hyperspectral image were first classified using the spectral information divergence method (Chein, 1999) and a binary mask was applied to segment out the remaining pixels (i.e. the plant pixels). Detected plant pixels were averaged to generate a representative reflectance spectrum with 256 spectral bands and resampled to a spectral width of 1 nm using a linear resampling approach (Lewitt, 1990). A total of 47 published VIs within the spectral range of 475 – 1710 nm was then computed (Table S1).

### Biochemical assays

Whole plant samples were finely ground using a pestle and mortar with liquid nitrogen, then subsampled separately for chlorophyll and sugar analysis and stored at –80 °C until biochemical analysis. For chlorophyll analysis, chlorophyll was extracted from 100 mg of sample with 100% methanol and centrifuged for 10 min at 10,016 g; this process was repeated twice. Extracts were analyzed by recording the absorbance at 750, 665, 652, and 470 nm using a UV-VIS spectrophotometer (Shimadzu UV-1800, Shimadzu Inc., Kyoto, Japan). Total chlorophyll was calculated using the formula described in Lichtenthaler (1987) and ranged between 396.2 – 821.9 μg/g in this study. For sugar analysis, soluble sugars were extracted from 100 mg samples with 80% ethanol and centrifuged for 10 min at 10,016 g; this process was repeated twice. Total soluble sugars were assayed according to the colorimetric method described in DuBois *et al*. (1956) and ranged between 2600 – 28300 μg/g in this study.

### Automated hyperspectral vegetation index (AutoVI) derivation

An automated system for hyperspectral vegetation index derivation named AutoVI, was developed to deliver effective VIs in a streamlined process covering model selection, model parameter generation, model parameter tuning, and model evaluation driven by a hyperparameter optimization framework, i.e. the optimizer, for plant phenotyping (Fig. 1). The optimizer is the core of the AutoVI method which seeks to generate a VI model for the desired trait based on a model evaluation metric, i.e. the objective function score (R^2^, see below) using an optimization algorithm given time and computing resource constraints. However, unlike simple optimization challenges which typically search for optimal solutions for a single model or function (static search space), multiple index models (dynamic search spaces) are optimized and evaluated in AutoVI (Fig. 1). This is made possible using dynamic parameter programming or “define-by-run” coding (Akiba *et al*., 2019), which generates the search space or set of model parameters during code execution depending on the index model or equation under evaluation.

**Figure 1.**
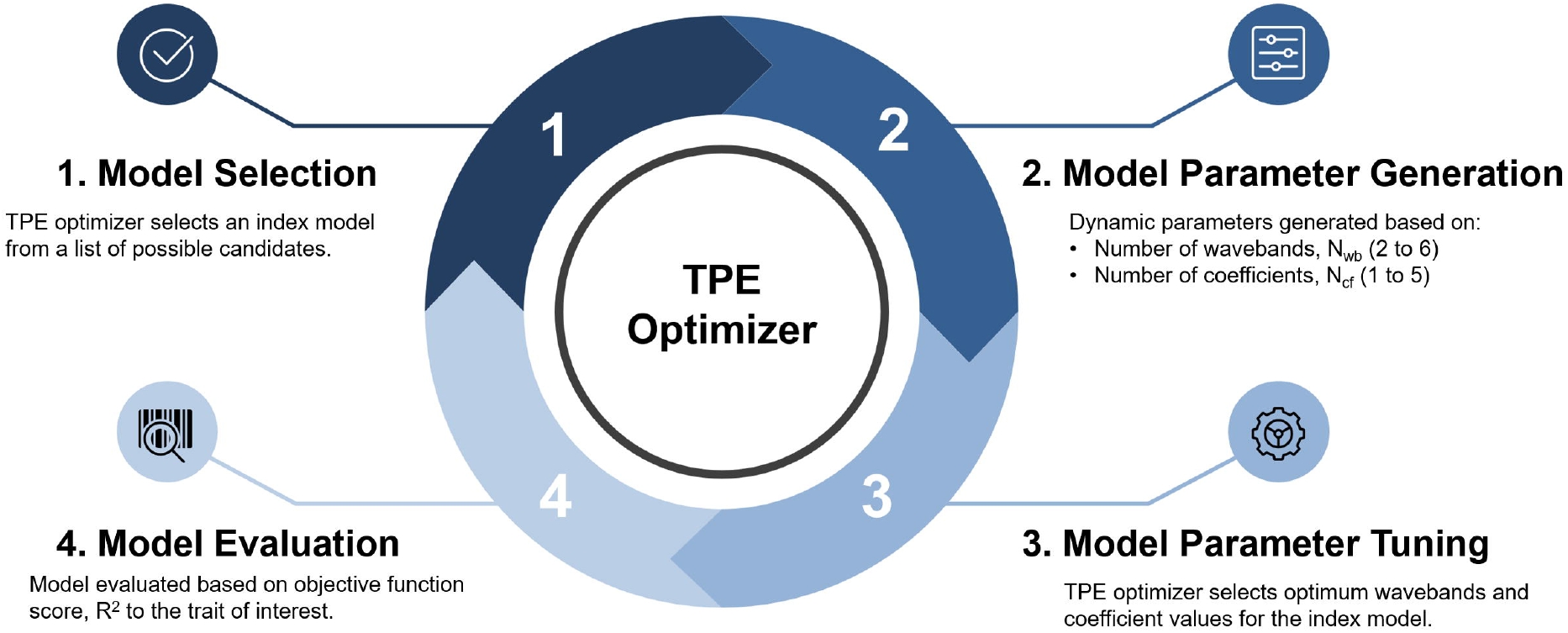
Overview of the AutoVI framework. The tree parzen estimator (TPE) algorithm is implemented as the optimizer for AutoVI. A single iteration consists of model selection, model parameter generation, model parameter tuning and model evaluation. AutoVI is programmed to repeat the optimization process until a pre-determined number of iterations is reached. The repeated iterations seek to maximize the objective function score, R^2^, by selecting the best candidate model and an optimum set of hyperparameters.

The mathematical expression (or model) defining a VI is primarily needed to combine two or more wavebands to decipher certain biochemical or biophysical traits of interest. The development of a new VI requires the selection of both a suitable model and a set of wavebands. A library of 33 index models was created (Table S2) after an exhaustive review of more than 500 previously developed VIs (Henrich *et al*., 2017). The listed model equations are of varying complexities, differing in the number of distinct wavebands (N_wb_ = 2 to 6, i.e. B1, B2,… B6) and the number of coefficients (N_Cf_ = 1 to 5, i.e. α, β,… ρ). The AutoVI system begins with model selection (step 1, Fig. 1) in which the optimizer selects a mathematical model at random from the library of 33 index models and generates the model parameters corresponding to the N_wb_ and N_Cf_ (step 2, Fig. 1). This is followed by model parameter tuning (step 3, Fig. 1) where optimum wavebands and coefficient values (between 0 – 1) are selected to maximize the objective function score (step 4, Fig. 1), which in this study is the coefficient of determination, R^2^, derived from a simple linear regression fitted to calculated index values and ground truth values for the measured trait of interest (in this case chlorophyll or sugar content). One run of optimization (steps I to 4) is referred to as a single iteration; the AutoVI system is programmed to repeat the optimization process until a pre-determined number of iterations is reached. At each iteration, an index model gets selected, and the computed objective function score (R^2^) is compared to the previous iteration. Additionally, unique sets of model parameters are computed for the respective index models. The repeated iterations seek to maximize the objective function score, R^2^, by selecting the best candidate model and an optimum set of model parameters.

In this study, the tree parzen estimator (TPE) (Bergstra *et al*., 2011; Yu & Zhu, 2020) was implemented as the optimization algorithm in AutoVI as it is widely employed for hyperparameter optimization in machine learning problems and showed better accuracy and efficiency compared to other algorithms when dealing with dynamic search spaces (Bergstra *et al*., 2015; Akiba *et al*., 2019; Yu & Zhu, 2020). The TPE algorithm is a variant of Bayesian optimization approaches which tries to construct a probabilistic model, also known as a “surrogate” model for mapping hyperparameters based on the probability of an objective function score given the set of hyperparameters. The AutoVI system, including the TPE algorithm customized to handle hyperspectral data was implemented in Python using the open source hyperparameter optimization library, Optuna version 2.0 (Akiba *et al*., 2019) with default settings. A graphical user interface (GUI) for AutoVI and source codes for TPE implementation are hosted on the public repository, GitHub (see Data availability). The AutoVI system was tested on an AMD Threadripper 3970X (32-cores) system with 256 GB RAM at the SmartSense iHub, Agriculture Victoria.

### AutoVI for chlorophyll content estimation

We evaluated the ability of AutoVI to derive high quality novel hyperspectral VIs for plant phenotyping using wheat’s total chlorophyll content as a biochemical trait. Chlorophyll content, either measured or estimated, can be a direct indicator of a plant’s primary production and has been used to determine N status and stress response of crop plants (Richardson *et al*., 2002; Murchie & Lawson, 2013). The existing dataset (n=80) was randomly split 65:35 into training and test datasets, with both datasets having the same sample distribution (i.e. stratified sampling) according to time points. The AutoVI system was trained on the training dataset to derive novel indices for chlorophyll content estimation, with the performance of these indices evaluated using the test dataset.

One possible issue with any optimization system is a selection bias towards index models with lower complexity, e.g. models with N_wb_ between 2 and 3 compared to those with higher complexity, e.g. models with N_wb_ ≥ 4. This is because the size of the solution search space increases exponentially with the increase in the number of input features i.e. N_wb_ (Winston, 1992; Yao & Liu, 1997). Consequently, more computational time or resource is required to optimize the complex models (N_wb_ ≥ 4) compared to the simpler models (N_wb_ ≤ 3) but their performance tend to scale better over time and do not plateau as fast as simpler models. When all model computations are grouped together, simpler models tend to outperform complex models in early stages of optimization, causing these to be favored by the optimizer and leading towards a ‘locally maximal solution’, which is the tendency of the computation to get stuck at a sub-optimal solution (Hinneburg & Keim, 1999). To address this issue in AutoVI, computations were performed on model groups consisting of models with the same N_wb_ (Fig. 2). Five parallel instances of AutoVI corresponding to the five model groups (M2, M3,… M6, equivalent to N_w_ = 2, 3,… 6) were executed with 20,000 iterations each with coefficients fixed at 1 (Fig. 2). These were repeated five times with the best performing index model from each group logged at each repetition (Fig. 2). This allows the reliable identification of the best index model for each model group and the overall best index model. The effect of longer optimizations and inclusion of coefficient on model performance was determined using the overall best index model, with five repeated AutoVI computations at 20,000 and 40,000 iterations with and without coefficient tuning.

**Figure 2.**
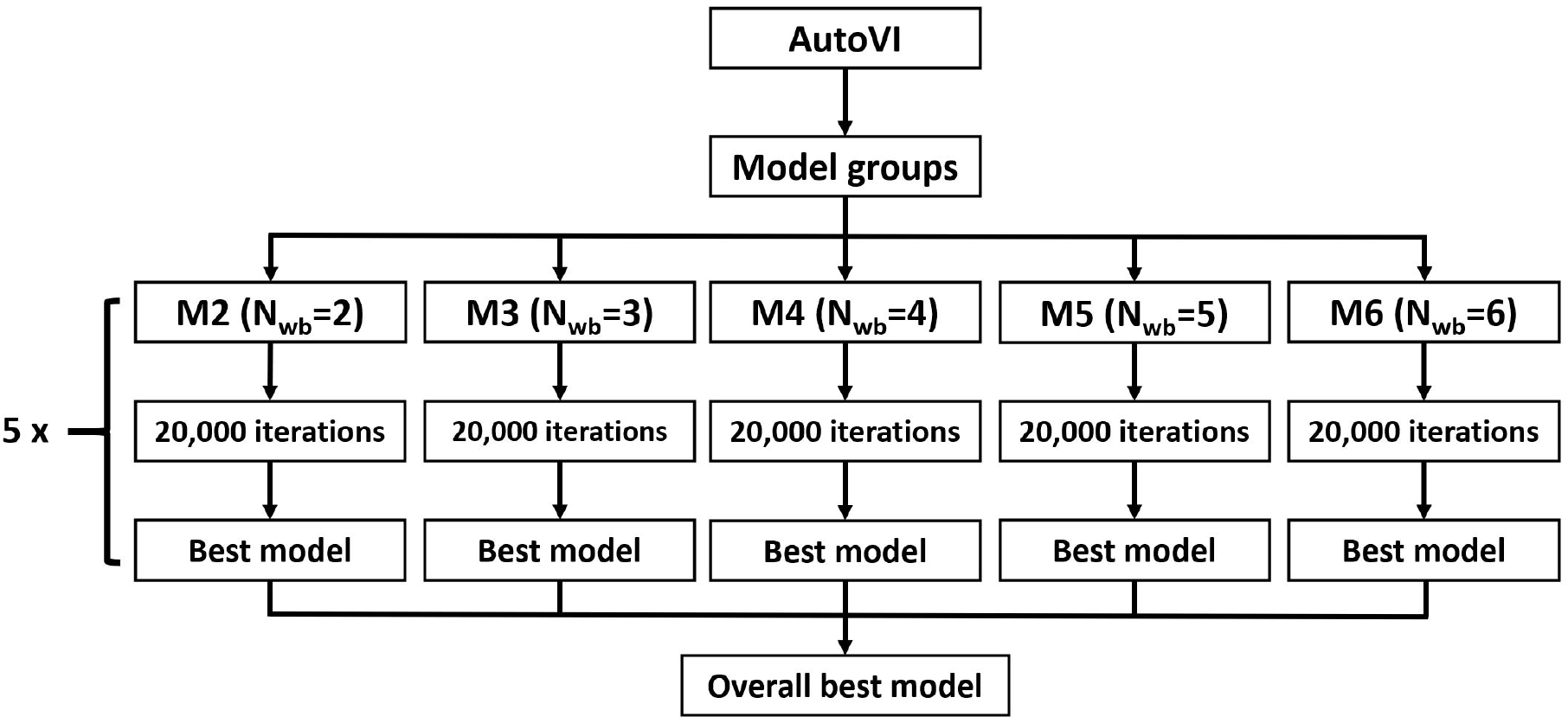
Flowchart for grouped model evaluations in AutoVI. Index models were grouped according to the number of wavebands (N_wb_) and parallel computations with 20,000 iterations each were performed on the model groups (M2, M3, M4, M5 and M6). The best index model for each model group and the overall best index model were identified from five repeated computations.

To determine the quality of AutoVI-derived indices for chlorophyll estimation, they were used as features in simple linear regression (SLR) modelling to predict chlorophyll content. SLR with each of the derived indices was first trained on the training dataset and then used to predict chlorophyll values for the test dataset. Model performance was evaluated using the R^2^ score calculated between predicted and actual chlorophyll values. In addition, performance for a stepwise multiple linear regression (SMLR) model (see below) with VIs selected from 25 AutoVI-derived indices (Fig. 2) is also included for comparison. Results achieved using AutoVI-indices were compared to those produced by SLR and SMLR with 47 published VIs, in addition to results provided by partial least-squares regression (PLSR) modelling using the full spectrum of reflectance values, i.e., reflectance values from all 1,235 wavebands (see below). For comparison across different regression models, additional performance metrics such as root mean squared error (RMSE), mean absolute error (MAE) and mean absolute percentage error (MAPE) are also provided.

### AutoVI for sugar content estimation

Sugar plays an important role in the osmotic adjustment of plants in response to drought stress. Studies have shown that genotypes that show higher accumulation of sugar content in leaves or stems are more drought tolerant (Adams *et al*., 2013; Piaskowski *et al*., 2016). Using the grouped model evaluation approach, AutoVI computations were conducted across five repetitions with 20,000 iterations each with the coefficients fixed at 1. The existing dataset was split 65:35 into training and test datasets as described previously, with the training of AutoVI conducted on the training dataset and validation of derived indices performed on the test dataset. Results for SLR and SMLR using AutoVI-derived indices were compared to those produced using 47 published VIs and PLSR modelling, as described in the previous section.

### Stepwise Multiple Linear Regression

Stepwise multiple linear regression (SMLR) is a feature selection method which iteratively adds (forward selection) or removes (backward selection) features to a multiple linear regression model to improve model performance, as indicated by an evaluation metric or score. In this study, we implemented a stepwise forward selection strategy based on a five-fold cross validated R^2^ score of a multiple linear regression model using the Python package, scikit-learn version 0.24. The maximum number of features to select was set to between 1 – 20 and selection was performed on 25 AutoVI-derived indices and 47 published VIs for both chlorophyll and sugar estimation on the training dataset. A multiple linear regression model was then fitted to the training dataset using the optimum selected features and used to predict target values (chlorophyll or sugar content) for the test dataset.

### Partial Least Squares Regression

Partial least-squares regression (PLSR) modelling is one of the most effective methods for plant trait prediction using hyperspectral data (Ely *et al*., 2019; Wu *et al*., 2019; Burnett *et al*., 2021). PLSR was designed to address both the collinearity between predictors, i.e., the different wavebands of a reflectance spectrum, and the large number of predictor variables when compared to trait observations. This study implemented PLSR modelling using the Python package, scikit-learn version 0.24 for chlorophyll and sugar content estimation. The optimal number of PLSR components was first determined based on five-fold cross validated R^2^ scores of PLSR models fitted on the training dataset with the number of components set to between 1 – 20 (Fig. S1). A PLSR model was then fitted to the training dataset using the optimum number of components (n=6 for chlorophyll and n=7 for sugar, Fig. S1) and used to predict target values (chlorophyll or sugar contents) for the test dataset.

## Results

### AutoVI for chlorophyll content estimation

AutoVI performance was measured using R^2^ scores generated by simple linear regression models on the test dataset with the respective AutoVI-derived indices as features. AutoVI performance across five repetitions was relatively stable, with the grouped evaluation strategy allowing for comparison across different model groups and identification of the best performing index model within each model group (Fig. 3). Between model groups, the M4 group (N_wb_ =4) had the best mean R^2^ of 0.7818, followed by M5 (N_wb_ =5) with R^2^ of 0.7747, M6 (N_wb_ =6) with R^2^ of 0.7689, M2 (N_wb_ =2) with R^2^ of 0.7637 and finally M3 (N_wb_ =3) with R^2^ of 0.7555. Overall, Index25 (N_wb_ =4, N_wb_=1) produced the best VI (R^2^=0.8007) and generated the best results across all five repetitions within the M4 group (Fig. 3). The performance of the best VIs according to model group is summarized in Table 1. Wavebands selected by AutoVI for the best chlorophyll indices derive predominantly from the red (600 – 720 nm) and red-edge (720 – 780 nm) regions, with a few wavebands from the blue (470 – 490 nm), near infrared (NIR, 1000 – 1300 nm) and shortwave infrared (SWIR, 1600 - 1700 nm) regions (Table 1). The R^2^ score produced by the best VI, termed hereafter as AutoVI chlorophyll index (AutoVI-Chl), was achieved using wavebands of 610 nm, 716 nm, 1384 nm and 1607 nm, as depicted in equation 1:

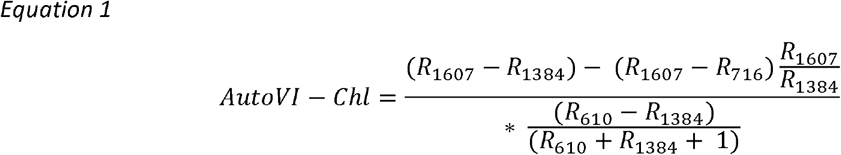

where, R_wb_ represent the reflectance measured at a discrete waveband (wb).

**Figure 3.**
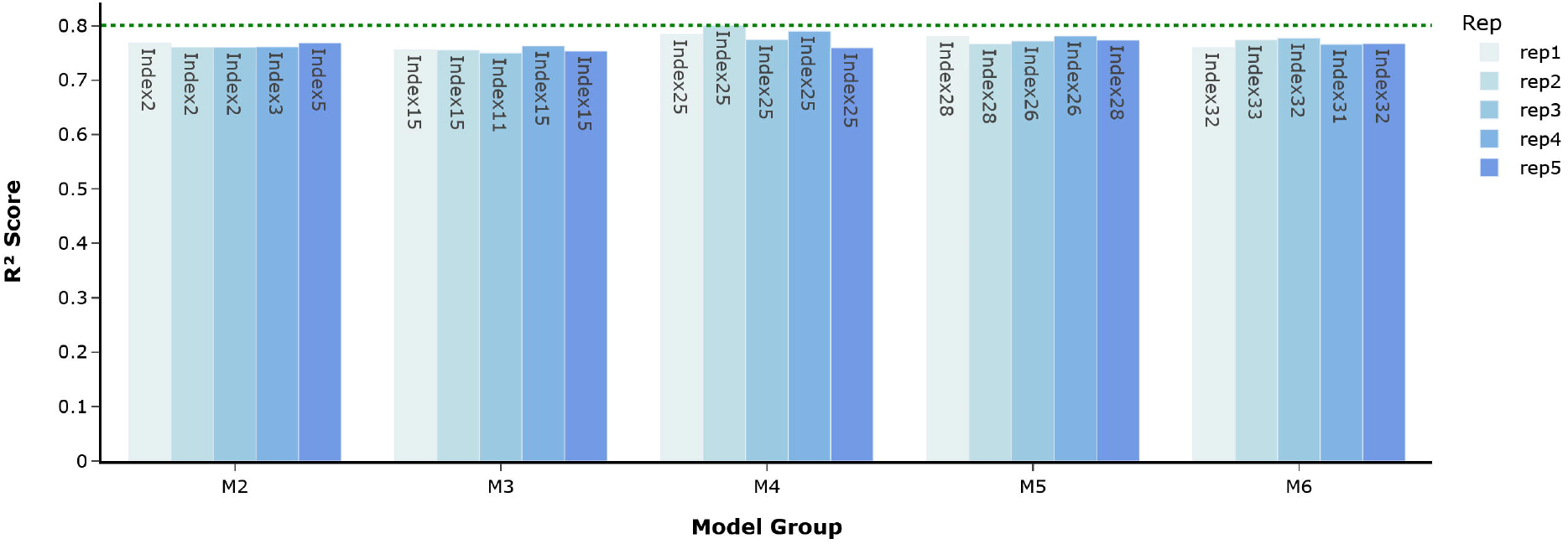
Best-performing index models from grouped model evaluations for total chlorophyll content estimation. The best index models from grouped (M2, M3, M4, M5 and M6) model evaluations were identified across five repeated AutoVI computations. Index model names are indicated within the bar figure. The overall best index model was Index25 from the M4 group with R^2^ score of 0.8007 (indicated by green dashed line).

**Table 1.** Performance of the best AutoVI-derived indices according to model group for chlorophyll estimation.

| Group | Index Model | Selected Wavelengths (nm) | R<sup>2</sup> | RMSE | MAE | MAPE |
| --- | --- | --- | --- | --- | --- | --- |
| M2 | 2 | 637, 711 | 0.7683 | 41.52 | 33.78 | 5.14% |
| M3 | 15 | 607, 716, 1712 | 0.7629 | 42.01 | 34.27 | 5.23% |
| M4 | 25 | 610, 716, 1384, 1607 | **0.8007** | **38.52** | **30.51** | **4.69%** |
| M5 | 26 | 497, 650, 707, 778, 1017 | 0.7813 | 40.35 | 32.74 | 5.13% |
| M6 | 32 | 477, 635, 713, 1033, 1035, 1668 | 0.7776 | 40.69 | 33.01 | 5.03% |

Performance metrics calculated for simple linear regression using the indicated AutoVI-derived index for chlorophyll estimation on the test dataset. The best scores are indicated in bold.

The performance of AutoVI-Chl with or without a coefficient variable, alpha (α), as depicted in its original equation (Index25, Table S2) was determined across five computational repetitions of 20,000 and 40,000 iterations (Fig. 4). At 20,000 iterations, the inclusion of the coefficient had minimal impact on AutoVI-Chl performance, as boxplots for R^2^ scores obtained with the coefficient (min=0.7657, median=0.7771, max=0.7959) and without the coefficient (min=0.7673, median=0.7875, max=0.7922) were comparable (Fig. 4). However, AutoVI computational time when the coefficient was included (∼2.1 hours for 5 repetitions) was up to 1.6x higher than without the coefficient (∼1.3 hours for 5 repetitions), suggesting that inclusion of coefficient(s) in AutoVI optimizations will likely incur computational costs. At 40,000 iterations, AutoVI-Chl performance deteriorated significantly with or without the coefficient (Fig. 4), suggesting that overfitting, where a model performs significantly better on the training (i.e. overfitted) but not the test dataset (i.e. unable to generalize to new data), is likely to occur with longer AutoVI runs. Based on these results, the recommended starting point for AutoVI training is up to 20,000 iterations and without coefficient tuning.

**Figure 4.**
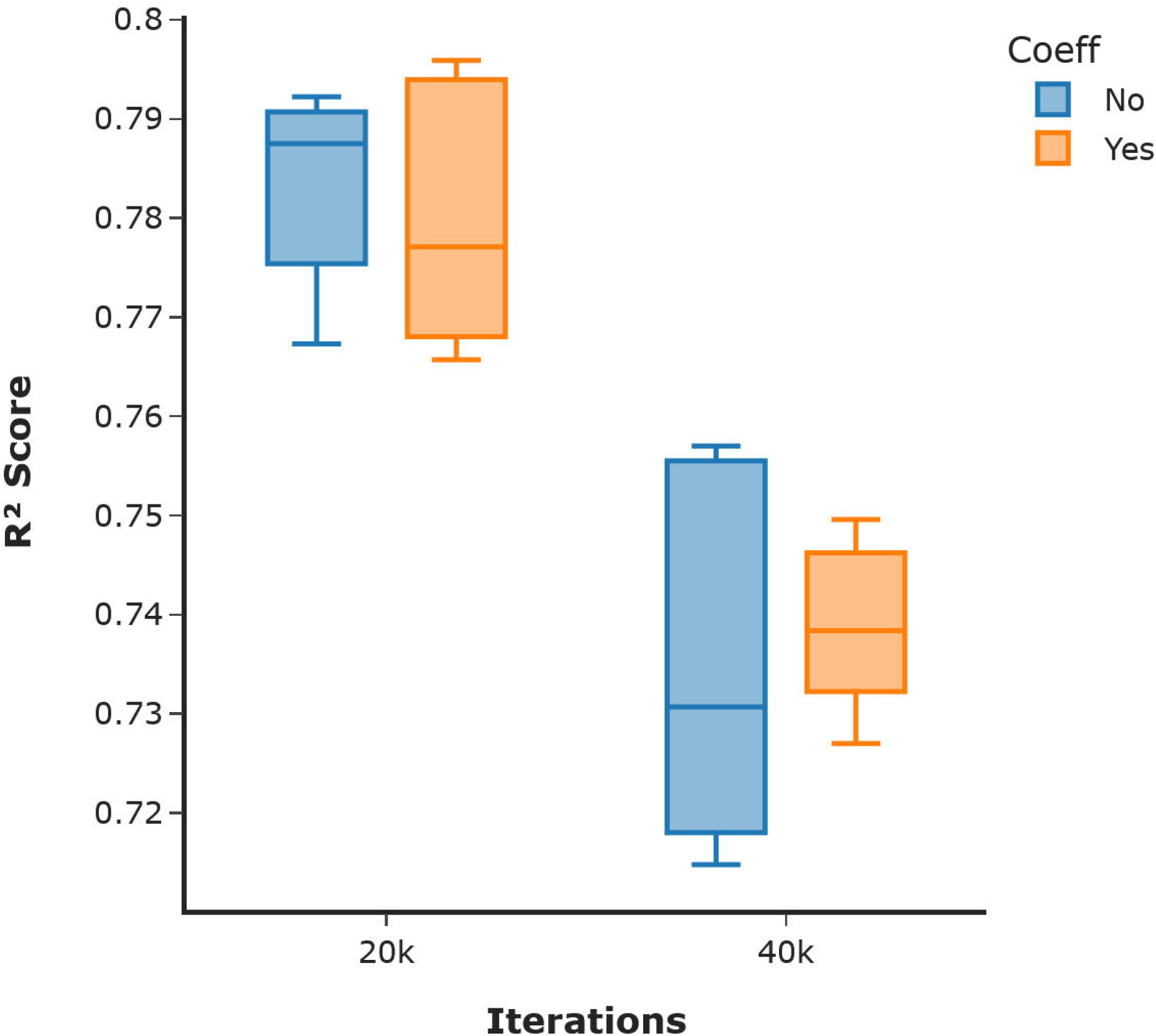
Effect of coefficient tuning and longer iterations on AutoVI chlorophyll index (AutoVI-Chl) performance. The inclusion (Yes) or exclusion (No) of a single coefficient, α, on AutoVI-Chl performance was determined across five repeated computations at 20,000 (20k) and 40,000 (40k) iterations each. The distribution of R^2^ scores are shown in the boxplot.

The quality of AutoVI-derived indices for chlorophyll content estimation was evaluated further against 47 published VIs, as features in simple linear regression (SLR) modelling. First, the best SLR model resulting from AutoVI-indices and the best SLR model with existing VIs was compared (Table 2). The model with AutoVI-Chl (R^2^=0.8007, RMSE=38.52, MAE=30.51, MAPE=4.69%) significantly outperformed the model with the normalized difference chlorophyll index, NDCI (R^2^=0.6018, RMSE=54.45, MAE=46.05, MAPE=7.09%) (Table 2, S3). Next, stepwise multiple linear regression (SMLR) models using the optimum subset of features selected from AutoVI-indices and existing VIs were compared (Table 2). For the existing VIs, SMLR with 7 VIs selected (Table S4) led to a significant improvement in model performance (R^2^=0.7136, RMSE=46.17, MAE=38.12, MAPE=5.88%) compared to SLR with NDCI but was still inferior to SLR with AutoVI-Chl; SMLR with four AutoVI-indices selected (Table S5) did not perform better (R^2^=0.7989, RMSE=38.69, MAE=30.98, MAPE=4.88%) compared to SLR with AutoVI-Chl. Finally, PLSR modelling performance for chlorophyll estimation was included as an additional benchmark for comparison. The PLSR model (R^2^=0.7379, RMSE=44.17, MAE=33.81, MAPE=5.36%) did not perform as well as SLR with AutoVI-Chl or SMLR with the selected AutoVI-indices (Table 2). Overall, the best modelling performance was provided by SLR with AutoVI-Chl. These results support AutoVI as an efficient system for novel trait-specific hyperspectral VI derivation.

**Table 2.**
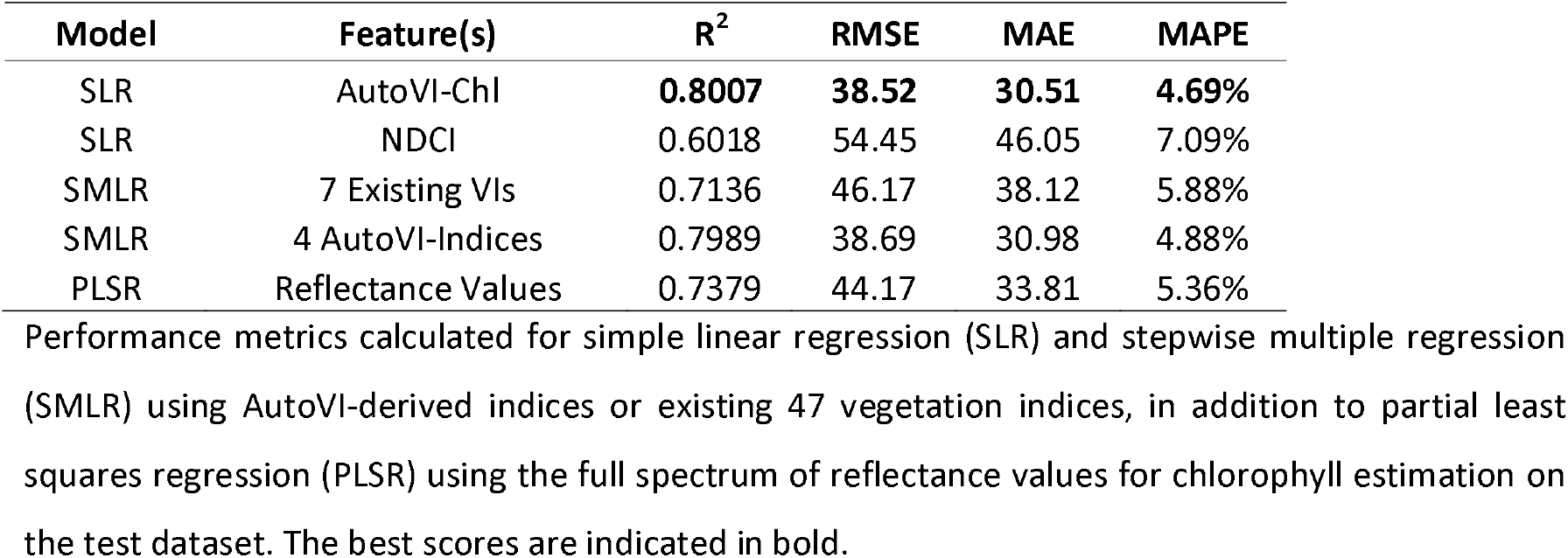
Comparison between different regression models for chlorophyll estimation.

### AutoVI for sugar content estimation

AutoVI performance across five repetitions was relatively stable, with the M6 group having the best mean R^2^ of 0.8201, followed by M3 group with R^2^ of 0.8127, M4 group with R^2^ of 0.7933, M5 group with R^2^ of 0.7877 and finally M2 group with R^2^ of 0.7591 (Fig. 5). The overall best VI was produced by Index33 (N_wb_=6, N_cf_=1), which also generated the best results for three repetitions within the M6 group (Fig. 5). The performance of the best VIs according to model group is summarized in Table 3. Wavebands selected by AutoVI for the best sugar indices derive predominantly from the shortwave infrared (1400 – 1700 nm) and near infrared (770 – 1370 nm) regions, with a few bands from the VIS (499 – 644 nm) region (Table 3). The R^2^ score produced by the best VI, termed hereafter as AutoVI sugar index (AutoVI-Sgr), was achieved using wavebands of 499 nm, 773 nm, 1179 nm, 1291 nm, 1425 nm and 1661 nm, as depicted in equation 2:

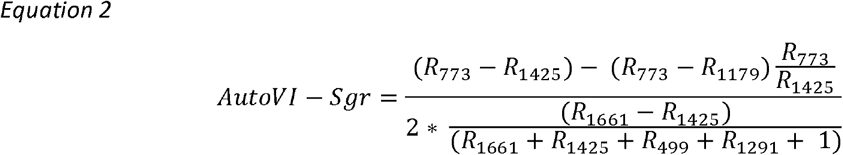

where, R_wb_ represent the reflectance measured at a discrete waveband (wb).

**Figure 5.**
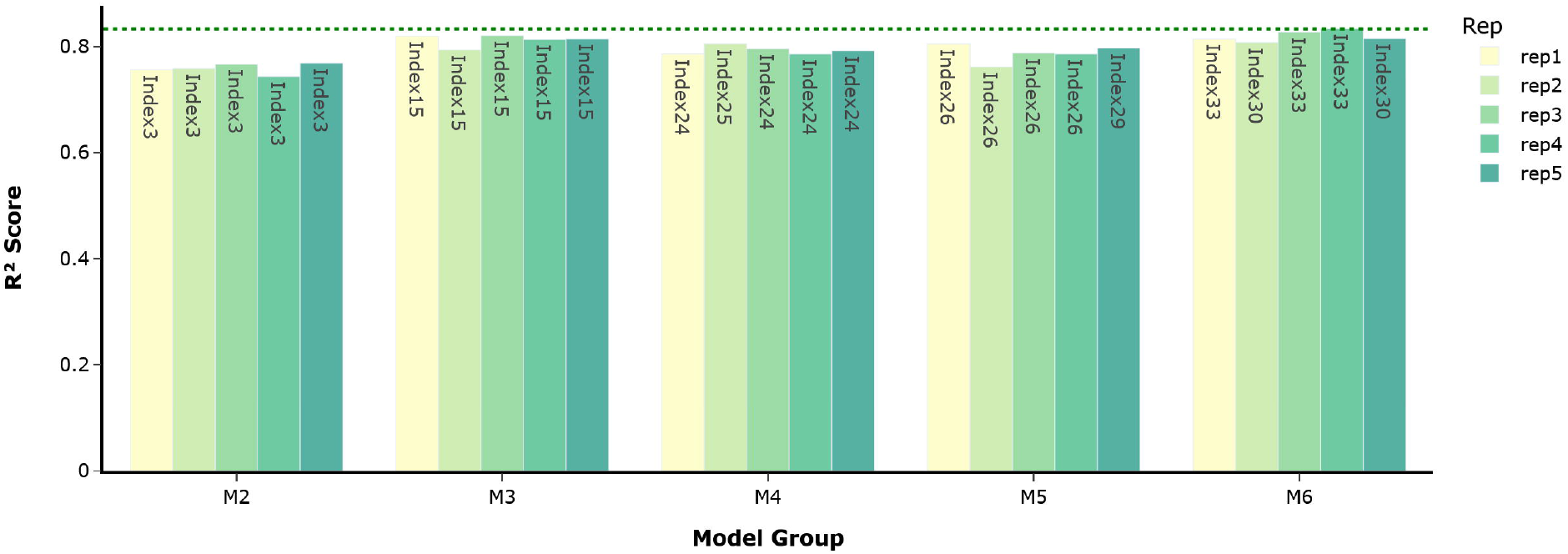
Best-performing index models from grouped model evaluations for total sugar content estimation. The best index models from grouped (M2, M3, M4, M5 and M6) model evaluations were identified across five repeated AutoVI computations. Index model names are indicated within the bar figure. The overall best index model was Index33 from the M6 group with R^2^ score of 0.8339 (indicated by green dashed line).

**Table 3.** Performance of the best AutoVI-derived indices according to model group for sugar estimation.

| Group | Index Model | Selected Wavelengths (nm) | R<sup>2</sup> | RMSE | MAE | MAPE |
| --- | --- | --- | --- | --- | --- | --- |
| M2 | 3 | 1420, 1588 | 0.7693 | 3078.89 | 2493.21 | 22.94% |
| M3 | 15 | 1271, 1417, 1650 | 0.8209 | 2712.72 | 2237.33 | 20.31% |
| M4 | 24 | 1371, 1422, 1650, 1712 | 0.8055 | 2827.07 | 2466.19 | 23.57% |
| M5 | 29 | 644, 990, 1406, 1418, 1649 | 0.7860 | 2964.88 | 2387.41 | 21.66% |
| M6 | 33 | 499, 773, 1179, 1291, 1425, 1661 | **0.8339** | **2612.41** | **2148.15** | **20.17%** |

Performance metrics calculated for simple linear regression using the indicated AutoVI-derived index for chlorophyll estimation on the test dataset. The best scores are indicated in bold.

The quality of AutoVI-derived indices for sugar content estimation was evaluated further against 47 published VIs, as features in simple linear regression (SLR) modelling. The best SLR performance with AutoVI-derived indices was achieved using AutoVI-Sgr (R^2^=0.8339, RMSE=2612.41, MAE=2148.15, MAPE=20.17%), which significantly outperformed the best SLR model achieved with the published VI, Gitelson and Merzlyak Index 2, GMI2 (R^2^=0.4695, RMSE=4668.35, MAE=3939.59, MAPE=38.96%). In general, SLR modelling performance with existing VIs was very poor (Table S6). SMLR with five existing VIs selected (Table S7) produced dramatically better results (R^2^=0.7387, RMSE=3276.50, MAE=2401.14, MAPE=22.30%) compared to SLR with GMI2 but was still inferior to the model with AutoVI-Sgr (Table 4). On the other hand, SMLR with four AutoVI-indices selected (Table S8) performed better (R^2^=0.8587, RMSE=2409.19, MAE=2071.47, MAPE=19.16%) compared to SLR with AutoVI-Sgr (Table 4). PLSR modelling performance for sugar estimation was also included as a benchmark for comparison. The PLSR model had similar performance (R^2^=0.8322, RMSE=2625.99, MAE=2212.89, MAPE=21.19%) as SLR with AutoVI-Sgr but was outperformed by SMLR with the AutoVI-indices (Table 4). These results further support AutoVI as an efficient system for novel VI derivation, with AutoVI-indices as high-quality features for trait prediction.

**Table 4.** Comparison between different regression models for sugar estimation.

| Model | Feature(s) | R<sup>2</sup> | RMSE | MAE | MAPE |
| --- | --- | --- | --- | --- | --- |
| SLR | AutoVI-Sgr | 0.8339 | 2612.41 | 2148.15 | 20.17% |
| SLR | GMI2 | 0.4695 | 4668.95 | 3939.59 | 38.96% |
| SMLR | 5 Existing VIs | 0.7387 | 3276.50 | 2401.14 | 22.30% |
| SMLR | 4 AutoVI-Indices | **0.8587** | **2409.19** | **2071.47** | **19.16%** |
| PLSR | Reflectance Values | 0.8322 | 2625.99 | 2212.89 | 21.19% |

Performance metrics calculated for simple linear regression (SLR) and stepwise multiple regression (SMLR) using AutoVI-derived indices or existing 47 vegetation indices, in addition to partial least squares regression (PLSR) using the full spectrum of reflectance values for sugar estimation on the test dataset. The best scores are indicated in bold.

## Discussion

In this study, we describe the design and implementation of an automated system for hyperspectral vegetation index derivation, AutoVI, for plant phenotyping using as examples chlorophyll and sugar content estimation in wheat. In both cases, indices generated by AutoVI significantly outperformed existing VIs in simple and multiple linear regression modelling, with performance exceeding that of a more complex model such as PLSR. The success of AutoVI can be attributed to the use of a highly performant hyperparameter optimization algorithm, tree parzen estimator (TPE) (Bergstra *et al*., 2011; Akiba *et al*., 2019).

Results in this study showed that AutoVI was able to efficiently generate high-quality 2- to 6-band indices specific to chlorophyll and sugar content. A review of existing literature highlights that most VIs currently deployed consist predominantly of 2-band indices, and to a lesser extent 3-band indices (Henrich *et al*., 2017). Indices with 4 bands or more are rare, presumably due to the difficulty in optimizing such indices (e.g. band selection) attributed to the curse of high dimensionality and issues associated with the multicollinearity of bands. Although a myriad of hyperspectral band selection strategy exists (Sun & Du, 2019), most of these were explicitly developed for image classification or regression models. Previous studies which focused on generating novel hyperspectral VIs used single or multiple correlation matrices between VI pairs and the trait of interest to uncover new band or index combinations (Thenkabail *et al*., 2004; Aasen *et al*., 2014; Xu *et al*., 2019). However, these approaches are computationally expensive as every possible combination of available bands (filtered or unfiltered) needs to be computed. Therefore, previous efforts have focused only on limited combinations between a few bands (2- and 3-band indices), thereby also limiting the net performance of the derived VIs. Consequently, the limited availability of specific VIs as biomarkers for plant traits has forced researchers to generalize upon the applicability of existing VIs, for example, the NDVI (2-band index), across almost all aspects of research and analysis, leading to suboptimal results. The AutoVI system is well-positioned to address this issue through the rapid generation of trait-specific novel 2-band to 6-band VIs without requiring band filtering or dimension reduction techniques to limit the number of input hyperspectral bands prior to processing.

Biochemical constituents in plants absorb electromagnetic energy in specific wavelength regions. Vegetation indices profiled around these characteristic spectral absorption regions can detect or estimate the biochemical trait of interest. The AutoVI system is constructed to automate the identification of these critical spectral regions using the underlying TPE algorithm. For chlorophyll, the selected wavebands centred around the red (600 – 700nm) and red-edge (700 – 740 nm) regions, with a few bands from the blue (470 – 490 nm), NIR (1000 – 1300 nm) and SWIR (1600 - 1700 nm) regions. Chlorophylls (chlorophyll *a* and *b*) are the most important plant pigments which function as photoreceptors and catalysts for photosynthesis, the photochemical synthesis of carbohydrates (Blackburn, 2006). As such, chlorophyll content in leaves and canopies is a key indicator of physiological measures such as photosynthetic capacity, developmental stage, productivity and stress (Richardson *et al*., 2002; Murchie & Lawson, 2013). Studies have shown that reflectance of wavelengths in the red region (∼530 – 630 nm, and a narrower band around 700 nm) is most sensitive to chlorophyll pigment concentrations across the normal range found in most leaves and canopies (Lichtenthaler *et al*., 1996; Gitelson *et al*., 2005). In addition, research has also shown that bands within the red-edge region (RE, 680 – 740 nm), which delineates the border between chlorophyll absorption in red wavelengths and leaf scattering in the NIR wavelengths, are strongly correlated with chlorophyll content (Curran *et al*., 1991; Gitelson *et al*., 1996). Existing chlorophyll VIs consist mainly of 2-band indices derived from ratios of narrow bands within regions of spectrum sensitive to chlorophyll pigments (VIS-RE, 400 – 740 nm) and those areas not sensitive to the pigments and/or related to some other control on reflectance (typically NIR, 750 – 900 nm) (Blackburn, 2006; Wu *et al*., 2008). Wavebands selected by AutoVI agree with published studies and additional wavebands selected from the SWIR region likely enhanced the sensitivity of some of the AutoVI-indices to chlorophyll by acting as control on reflectance.

The sugar metabolic pathway is intrinsically linked with the regulation of plant growth and development, and response to stress (Julius *et al*., 2017; Kaur *et al*., 2021). Studies have shown that abiotic stress such as drought or heat triggers sugar accumulation, particularly soluble sugars such as sucrose in plants (Lemoine *et al*., 2013; Zhou *et al*., 2017). In this study, AutoVI generated novel indices with a strong correlation to total sugar content, with specific wavebands selected mainly from the SWIR (1400 – 1700 nm) and NIR (770 – 1370 nm) regions. It is known that leaf reflectance properties in the SWIR region (1300 – 2500 nm) is governed by water content and biochemical compounds such as cellulose, sugars and starch (Elvidge, 1990; Kokaly *et al*., 2009). Indeed, more recent studies have used near infrared spectroscopy (750 – 2500 nm) coupled with machine learning and/or statistical models to estimate soluble carbohydrates, including total sugar in various plant tissue and organ such as leaf and stem (Adams *et al*., 2013; Piaskowski *et al*., 2016). Research in rice has also identified wavebands in the NIR region (800 – 1100 nm) as being important for sugar content estimation in leaves (Das *et al*., 2018). These studies provide support for bands selected by AutoVI as being specific to total sugar content. Models for sugar estimation in this study were less robust overall (e.g. MAPE = 19.16 – 22.30%) compared to models for chlorophyll estimation (e.g. MAPE = 4.69 – 5.23%) as reflected in the higher error scores. This may be due to difficulties in obtaining high quality spectral signatures from the SWIR region as water vapor is known to obscure spectral signatures for biochemical compounds in this region in plants (Elvidge, 1990; Kokaly *et al*., 2009). Nevertheless, it is noteworthy that AutoVI was able to select bands in the SWIR region whilst avoiding the water absorption peak at 1450 nm (Elvidge, 1990; Kokaly *et al*., 2009).

Novel AutoVI-derived indices for chlorophyll and sugar content estimation were first compared against 47 published VIs as features in simple linear regression (SLR). This provided a good baseline to compare VIs as model performance would depend solely on the quality of the VI. For chlorophyll estimation, the best SLR achieved with the AutoVI-derived index, AutoVI-Chl (R^2^=0.8007) significantly outperformed the best SLR with the published VI, NDCI (R^2^=0.6018). Similarly, for sugar estimation, the best performance provided by SLR with AutoVI-Sgr (R^2^=0.8339) significantly outperformed the SLR with the published VI, GMI2 (R^2^=0.4695). Next, the performance of a stepwise multiple linear regression (SMLR) with features selected from existing VIs was compared to the SLR models with AutoVI-indices. Results showed that although SMLR did significantly improve performance for both chlorophyll (R^2^=0.7136) and sugar (R^2^=0.7387) estimation with published VIs, these were still inferior compared to SLR results achieved using the AutoVI-indices. SMLR was also performed using the AutoVI-indices, with performance enhancement observed for sugar (R^2^=0.8587) but not chlorophyll (R^2^=0.7989) estimation. Finally, model performance for chlorophyll and sugar estimation using AutoVI-indices were compared to results produced by PLSR, a well-established method for plant trait modelling using hyperspectral data (Ely *et al*., 2019; Wu *et al*., 2019; Burnett *et al*., 2021). PLSR represents a different modelling approach as it projects the entire spectrum of reflectance values (or the predictor variables) into a smaller number of variables (or components) whilst simultaneously maximizes the correlation between the response and the variables (Geladi & Kowalski, 1986; Wold *et al*., 2001). With PLSR, what is being compared to the models with AutoVI-indices is the quality of feature representation or learning as depicted by the projected components in PLSR in contrast to the functions encoded in the AutoVI-indices. For both chlorophyll and sugar estimation, SLR and SMLR with AutoVI-indices outperformed PLSR (R^2^=0.7379 for chlorophyll; R^2^=0.8322 for sugar). Impressively, SLR with the best AutoVI-derived index (AutoVI-Chl or AutoVI-Sgr) produced better results and outperformed more complex approaches such as SMLR (except for sugar estimation) and PLSR. These results provide strong support for AutoVI-derived indices as high-quality features and affirm AutoVI as a high-performing system for novel trait-specific hyperspectral VI derivation.

However, AutoVI-indices, including AutoVI-Chl and AutoVI-Sgr, are not perfected VIs and will benefit from further studies incorporating more data collected from multiple genotypes across different environments and growth stages. Particularly for field-based phenotyping, AutoVI-indices should ideally be derived using canopy spectra, as these are known to differ from leaf spectra (Croft *et al*., 2014). Depending on the underlying trait, it may be worthwhile to collect samples from different plant tissues or organ. For example, studies have shown that genotypes which accumulate more sugar in leaf and stem tissues are more heat and/or drought tolerant (Piaskowski *et al*., 2016; Zhou *et al*., 2017). Fortunately, AutoVI does not impose any size or dimensional constraint on the input data and is expected to work with data derived from different hyperspectral sensors and/or spectra sources. Depending on the data provided, AutoVI can be customized to deliver trait-specific VIs according to species, genotype, growth stage and environment, making it a powerful and versatile tool for both novice and expert users alike. In addition, compared to machine learning and statistical modelling approaches where model optimization and deployment remains technically challenging, AutoVI-derived indices can be easily computed and readily deployed for HTP without requiring complex hardware or software resources.

In conclusion, results in our study demonstrate that AutoVI is an efficient and powerful tool for deriving high-quality hyperspectral VIs for plant phenotyping. AutoVI-derived VIs outperformed existing VIs and delivered strong performance in SLR and SMLR modelling for chlorophyll and sugar content estimation, producing results superior to PLSR. The AutoVI system is expected to accelerate the development of novel VIs for plant/crop traits which will find wide application in HTP and agriculture remote sensing vital to improving breeding program and crop management efficiencies.

## Author contributions

JCOK and BPB designed the AutoVI system and analyzed the data. JCOK implemented the AutoVI system and GUI version in Python. BPB researched and collated the list of index models used in AutoVI. JCOK, BPB, GS and SK conceived and designed the study. JCOK, BPB, GS and SK wrote and edited the paper. All authors read and approved the final manuscript.

## Data availability

All relevant source codes and datasets, including a GUI implementation of AutoVI in Python for Windows 10 64-bit operating system required to reproduce results reported in this study are available at www.github.com/AVR-SmartSense/AutoVI.

## Supporting Information

**Figure S1. Optimization of number of components in PLSR for chlorophyll and sugar estimation. Table S1. List of 47 published vegetation indices**.

**Table S2. Index models for AutoVI selection**.

**Table S3. Performance of 47 published vegetation indices for chlorophyll content estimation**.

**Table S4. Stepwise multiple linear regression with 47 published vegetation indices for chlorophyll estimation**

**Table S5. Stepwise multiple linear regression with AutoVI-derived indices for chlorophyll estimation**

**Table S6. Performance of 47 published vegetation indices for sugar content estimation**.

**Table S7. Stepwise multiple linear regression with 47 published vegetation indices for sugar estimation**

**Table S8. Stepwise multiple linear regression with AutoVI-derived indices for sugar estimation**

