## Supplementary figures and images for "Automated hyperspectral vegetation index derivation using a hyperparameter optimization framework for high-throughput plant phenotyping"

### Supplemental Figure 1

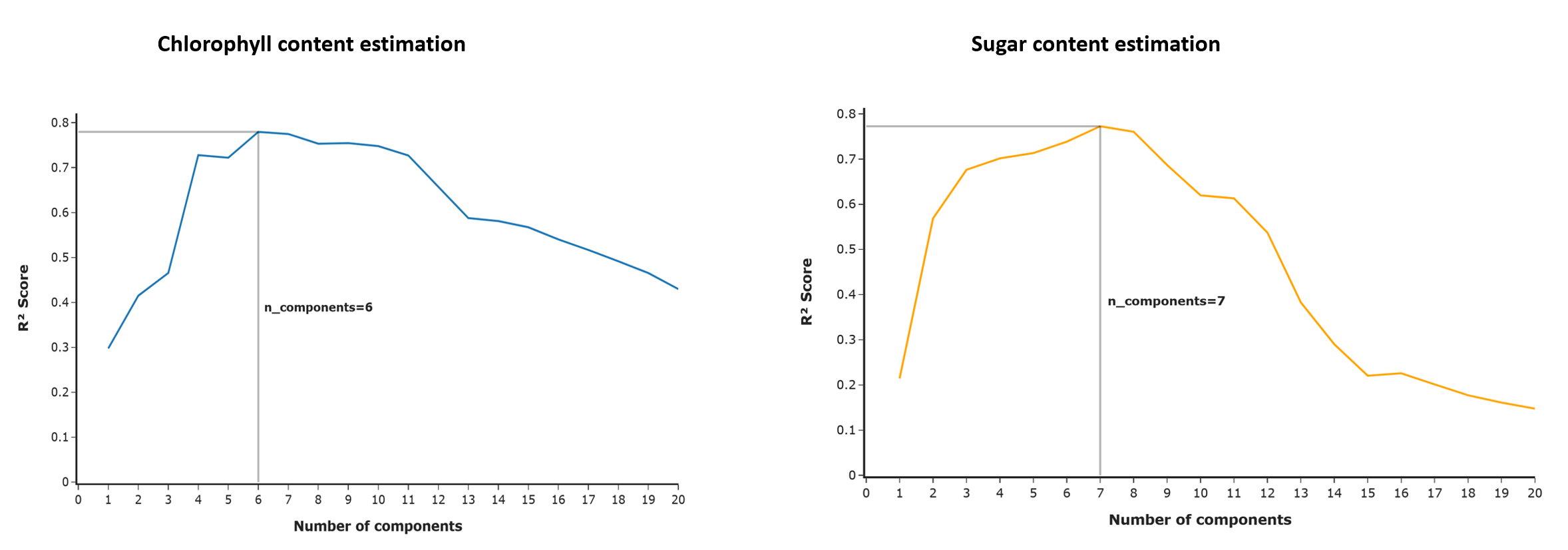
